# Poverty shapes the transcriptome of immune cells

**DOI:** 10.1101/2022.12.06.517536

**Authors:** Nicole S. Arnold, Justyna Resztak, David Witonsky, Adnan Alazizi, Nicole Noren Hooten, Michele K. Evans, Valerie Odero-Marah, Douglas F. Dluzen, Roger Pique-Regi, Francesca Luca

## Abstract

Social factors influence health outcomes and life expectancy. Individuals living in poverty often have adverse health outcomes related to chronic inflammation that affect the cardiovascular, renal, and pulmonary systems. Negative psychosocial experiences are associated with transcriptional changes in genes associated with complex traits. However, the underlying molecular mechanisms by which poverty increases the risk of disease and health disparities are still not fully understood. To bridge the gap in our understanding of the link between living in poverty and adverse health outcomes, we performed RNA sequencing of blood immune cells from 204 participants of the Healthy Aging in Neighborhoods of Diversity across the Life Span (HANDLS) study in Baltimore, Maryland. We identified 138 genes differentially expressed in association with poverty. Genes differentially expressed were enriched in wound healing and coagulation processes. Of the genes differentially expressed in individuals living in poverty, *EEF1DP7* and *VIL1* are also associated with hypertension in transcriptome-wide association studies. Our results suggest that living in poverty influences inflammation and the risk for cardiovascular disease through gene expression changes in immune cells.

## Introduction

Socioeconomic disparities in health and mortality are very well documented^1–3^. These inequalities have been observed at similar levels since the early nineteenth century. More recently, health disparities are also reflected in current major causes of death including cancer and cardiovascular disease^4^. Individuals living in poverty are more frequently exposed to risk factors for these conditions including poor diet, inadequate exercise, and smoking. Additionally, psychosocial factors that may influence health status are also common when living in poverty, such as housing insecurity, discrimination, structural racism, and lack of social support ^5^. Living below the federal poverty line increases cardiovascular risk^6^ and low socioeconomic status (SES) is associated with an increase of cardiovascular disease biomarkers in the circulation, including red-blood-cell width^7^, high-density lipoprotein cholesterol (HDL-C)^8^, intimal medial thickness of the arteries and pulse-wave velocity^9^. These risks are also further amplified in vulnerable groups such as African American adults^9^. Low SES is associated with high prevalence of Cytomegalovirus seropositivity, which is also associated with higher chronic conditions^10^. Low SES has consistently been associated with a greater risk of infectious and cardiovascular diseases, more rapid progression, and decreased survival^4^. The biological mechanisms underlying these disparities still need to be better understood. Elevated innate immunity responsiveness is observed in low SES children when compared to children that live in a higher SES^11^. In a broad cohort across the United States, low childhood SES was linked to poorer metabolic outcomes in midlife^12^.

Adverse health is consistently linked to lower SES, but little is known about the molecular mechanisms behind this disparity. Individuals from low-income families have higher cortisol output in daily life, higher expression of transcripts containing NFkappaB response components, and higher production of the proinflammatory cytokine interleukin 6^13^. Children from low-income homes have increased cortisol levels in adulthood, regardless of social standing^13,14^, which could potentially lead to resistance to glucocorticoid signaling and allow for increased adrenocortical and inflammatory responses^13^. Additionally, metabolic syndrome components such as central obesity, insulin resistance, and blood pressure can worsen with chronic cortisol exposure^15^.

Social genomics investigates the various dimensions of a person’s social and psychological environment that influence gene expression^16–20^. Several studies have linked changes in gene expression in whole blood to socioeconomic status, psychosocial experiences, and adverse social conditions^21–27,28,29^. Earlier studies in the field^16,17,19^ identified a pattern of differentially expressed genes known as the conserved transcriptional response to adversity (CTRA) that are altered in response to various aspects of social adversity. The CTRA comprises genes in Inflammatory pathways as well as innate antiviral response and antibody production genes^30,31^. However, socioeconomic status and psychosocial experiences have a broader effect on immune cell gene expression and impact additional pathways beyond those included in the CTRA^28^.

Therefore, there is a need in the field to include genome-wide transcriptomic studies of individuals living in poverty and in a sample balanced by age, race, and sex. In a recent analysis of a cohort of 1,069 young adults from the National Longitudinal Study of Adolescent to Adult Health (Add Health) found gene expression changes with demographic and behavioral factors; yet family poverty was not found to be associated with individual genes unless aggregated by functional groups^32^. Characterizing whether living in poverty induces genome-wide transcriptional changes thus remains essential to understand the biological mechanisms underlying poor health outcomes and health disparities in immune and cardiovascular diseases among individuals living in poverty. Here we hypothesized that differential gene expression is associated with poverty status and that understanding the molecular pathways affected by poverty may shed light on health disparities in disease risk and outcome. We analyzed a sub-cohort of participants from the National Institute on Aging’s (NIA) Healthy Aging in Neighborhoods of Diversity Across the Life Span (HANDLS) study in Baltimore, MD. To determine the association between poverty status and gene expression, we studied 204 participants from the HANDLS cohort and performed RNA-sequencing on peripheral blood mononuclear cells (PBMCs) collected at two different time points (Figure 1).

**Figure 1:**
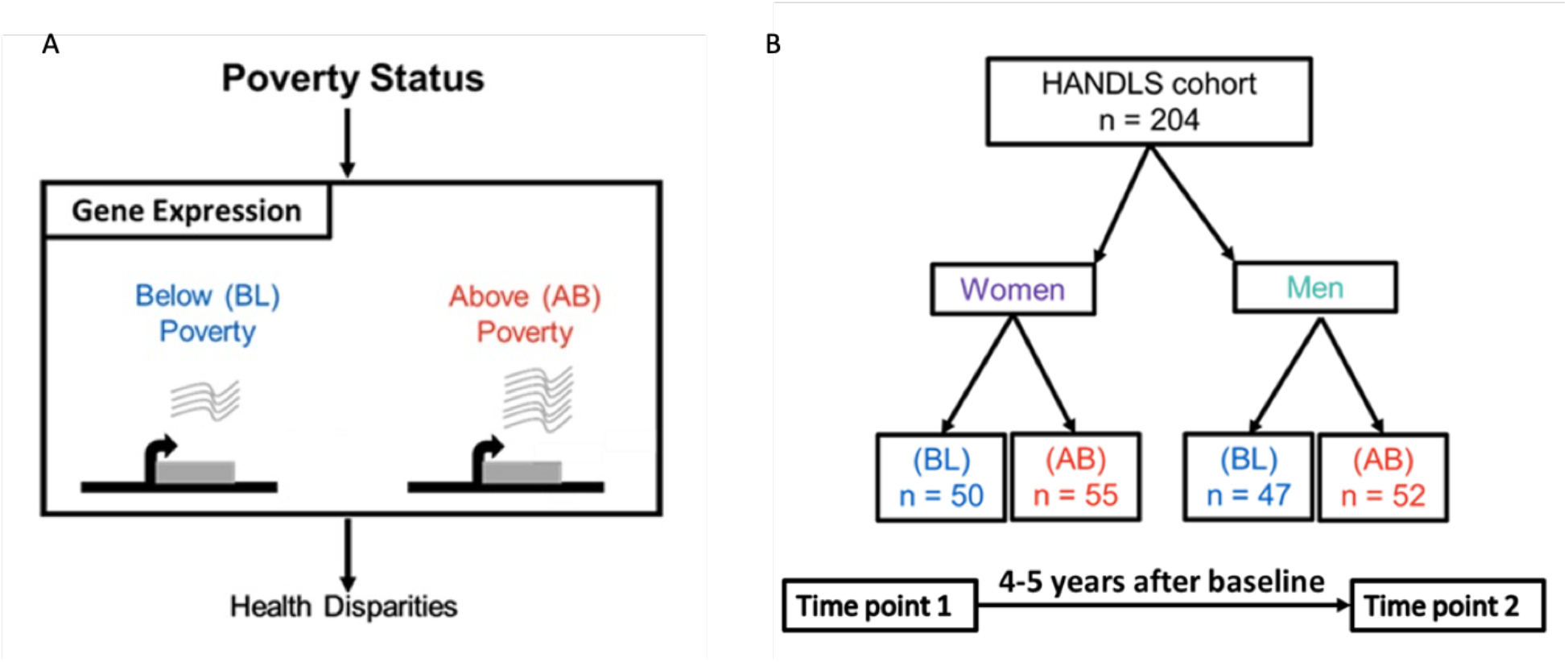
Study Design. A. Conceptual framework showing the workflow of the study to examine the relationship between poverty status and gene expression patterns leading to health disparities. Below the poverty line *BL* is shown in blue, and above the poverty line *AB* is shown in red. B. Study cohort. The final sample cohort consists of 204 participants. Blood samples were collected at two different time points ∼ 4-5 years apart.

## Results

### Poverty status influences gene expression in the immune system

To identify genes differentially expressed between individuals living in poverty and those living above poverty, we analyzed differential gene expression using DESeq2 at each time point. We controlled for technical variables, sex, age, and the first three genotype PCs (Supplementary Figure 1). We observed 138 differentially expressed genes at the first time point (FDR<10%). We found that 70% of the differentially expressed genes had lower expression in individuals living in poverty (Figure 2B).

**Figure 2:**
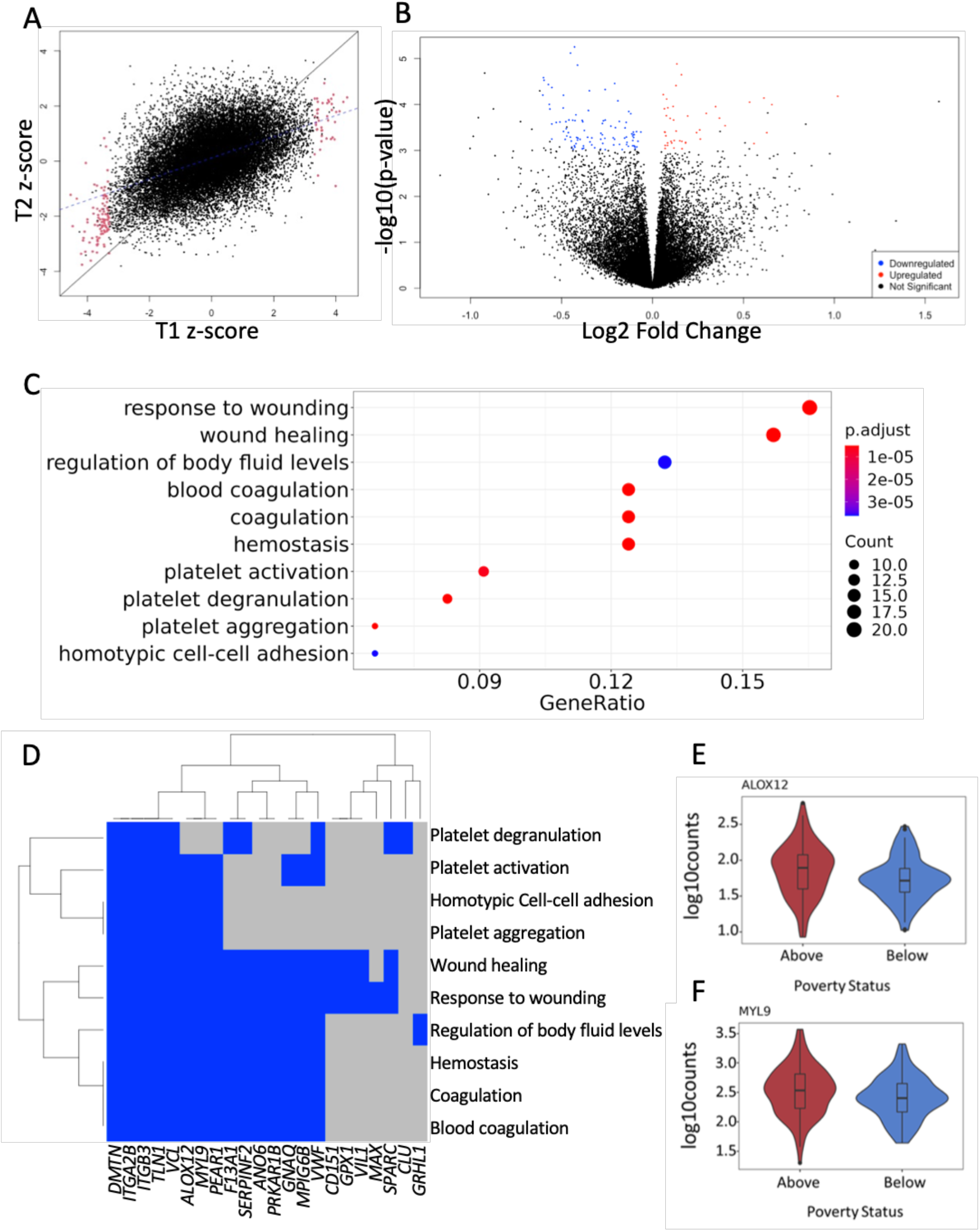
Poverty status influences gene expression in immune cells. A. Scatter plot of the correlation between gene expression changes associated with poverty (z-scores) in each time point separately. Significantly differentially expressed genes at time point 1 are indicated in red (FDR<10%). B. Volcano plot of the differentially expressed genes at time point 1. The 96 downregulated genes are indicated in blue and the 42 upregulated genes are in red. C. Results of the enrichment analysis for the gene ontology terms in the differentially expressed genes. The x-axis displays the proportion of gene (gene ratio), y-axis top ten enriched gene ontology terms (biological processes), legend indicates the number of associated genes (n=10-20), and the p.adjust legend. D. Heatmap of significantly differentially expressed genes in the top 10 enriched gene ontology terms. Blue indicates that a gene is down-regulated in individuals living in poverty, grey indicates that the gene is not part of the gene ontology term. E-F. Examples of differentially expressed genes. The x-axis indicates the poverty status (as in Figure 1), the y-axis indicates the normalized log10 expression counts.

We wanted to validate these findings further by also comparing differentially expressed genes from time point one to time point two. We found that 74 of the 138 genes are differentially expressed at the second time point (p<0.05), and overall we find consistent effects between the two time points as indicated by a strong correlation in the gene expression z-score (r=0.47, p<10^−16^) (Figure 2A). This result confirms that the gene expression changes associated with poverty are largely consistent across time points. We then asked whether the gene expression changes associated with poverty are specific to our cohort or shared with an independent cohort of 1,069 young adults from the National Longitudinal Study of Adolescent to Adult Health (Add Health)^32^, where the effect of poverty on gene expression was also tested. We found a significant correlation between the z-scores from the two studies (ρ=0.264 and p-value< 2.2e^-16^, Supplementary Figure 2).

We performed an over-representation analysis to identify the biological processes impacted by poverty. We found an enrichment for biological processes pertaining to wound healing, blood coagulation, and platelet activation (Figure 2C-D) with largely overlapping sets of differentially expressed genes in these biological categories. Among these genes, *ALOX12* (log2 fold change (LFC)= -0.57, p-adj < 0.06) and *MYL9* (LFC = -0.53, p-adj < 0.09) have the largest absolute change in expression and are in several enriched GO categories, including response to wounding, blood coagulation, and platelet aggregation (Figure 2E-F, Supplemental Table 3). *ALOX12 (*Arachidonate 12-Lipoxygenase, 12S Type) encodes lipid metabolizing enzymes that regulate inflammation, platelet aggregation, cellular migration, and modulation of vascular permeability^33^. Depletion of the gene *ALOX12* has been associated with chronic inflammation via effects on pro-inflammatory activity^33^. Direct evidence links *ALOX12* to chronic diseases characterized by unregulated inflammation^34,35^). Promoter methylation changes at *ALOX12* result in potential epigenetic alterations in the etiology of atherosclerosis plaques^36^. *ALOX12* is expressed in various cell types and organs throughout the body, linking to several diseases, including atherosclerosis, hypertension, diabetes, obesity, and neurodegenerative disorders^37^. Genetic variants in the *ALOX12* gene are associated with myocardial infarction, increased cardiovascular events, and mortality^38^. Myosin-9 *MYL9* is a motor protein that plays a role in cellular movement and contraction^39,40^.

### Effects of poverty on gene expression in men and women

Sex differences in physiology and pathology, along with sex steroid receptor signaling have an influence on health outcomes and contribute to life expectancy differences between men and women^41–43^. Sex hormones modulate the inflammatory system, and sexual dimorphism drives inflammatory activity^44,45^. It has been shown that sex influences gene expression levels and cellular composition of tissue samples across the human body, including immune cells^46^. To investigate whether transcriptional differences associated with poverty may affect men and women differently, we split our study cohort by sex and investigated gene expression patterns associated with poverty in men and women separately. We found 104 differentially expressed genes in women, 60 upregulated and 44 downregulated with poverty (Figure 3B). However, we did not observe any genes differentially expressed in men living in poverty (Figure 3A). This is also reflected in the observation that a larger proportion of genes have high z-scores in women (16.2% of tested genes with |z|>2), compared to men (3.6% of tested genes with |z|>2). This difference may be due to unmeasured confounding factors in men, or to a stronger effect of poverty on gene expression in immune cells in women.

**Figure 3:**
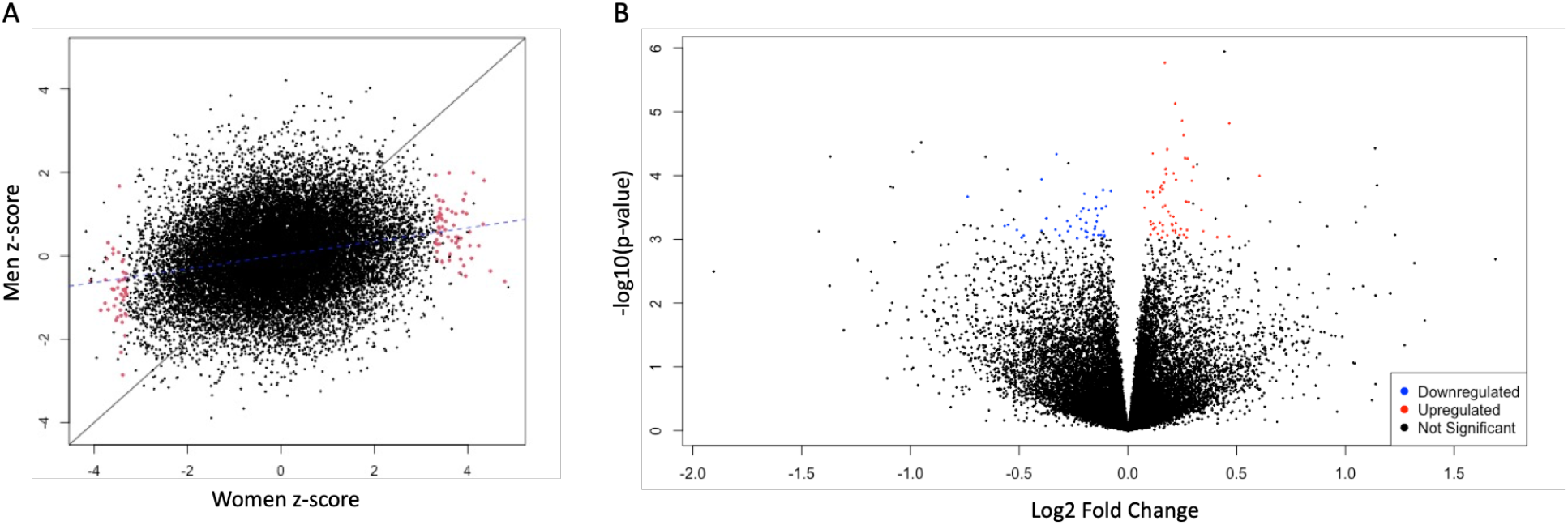
Association between poverty and gene expression in men and women. A. Scatter plot of the correlation between gene expression z-scores of men and women. Significantly differentially expressed genes are indicated in red (FDR<10%). B. Volcano plot of the differentially expressed genes in women. The 44 downregulated genes are indicated in blue and the 60 upregulated genes are in red.

### Understanding diseases and conditions influenced by poverty through transcriptional changes

To examine the adverse effects of poverty on disease risk that may be mediated by changes in gene expression in immune cells (Figure 1A), we considered genes associated with disease risk through a probabilistic transcriptome-wide association study (PTWAS^47^). Because of the enrichment in genes associated with coagulation and wound healing and the importance of inflammation in cardiovascular health, we focused on cardiovascular disease. We considered all genes significantly differentially expressed at time point 1 and found two genes associated with hypertension (Figure 4). Lower expression of *EEF1DP7* and *VIL1* was associated with increased risk for hypertension in PTWAS (Figure 4), and we also found lower expression of these genes in individuals living in poverty compared to those living above poverty, thus suggesting that poverty may impact the molecular pathways associated with hypertension. Furthermore, poverty status and disease risk both correlate in the same direction with gene expression, thus showing concordant^48^ effects. Note that the direction of the PTWAS effect also reflects the original context of the eQTL study and the LD structure in the discovery sample does not capture epistasis and co-regulation, therefore caution should be exercised in over-interpreting the sign of the z-score. *VIL1* encodes calcium-regulated actin-binding proteins that cap, sever, and build actin filaments^49,50^,^50^,^51^. In addition, *VIL1* encodes for the tissue-specific protein villin expressed in renal and gastrointestinal epithelial cells^52^. *VIL1* (LFC= -0.51, p-adj = 0.07) is in the pathways for response and regulation of wound healing. Lower expression of *VIL1* is associated with prolonged stress and ineffective recovery of cell stress^53^.

**Figure 4:**
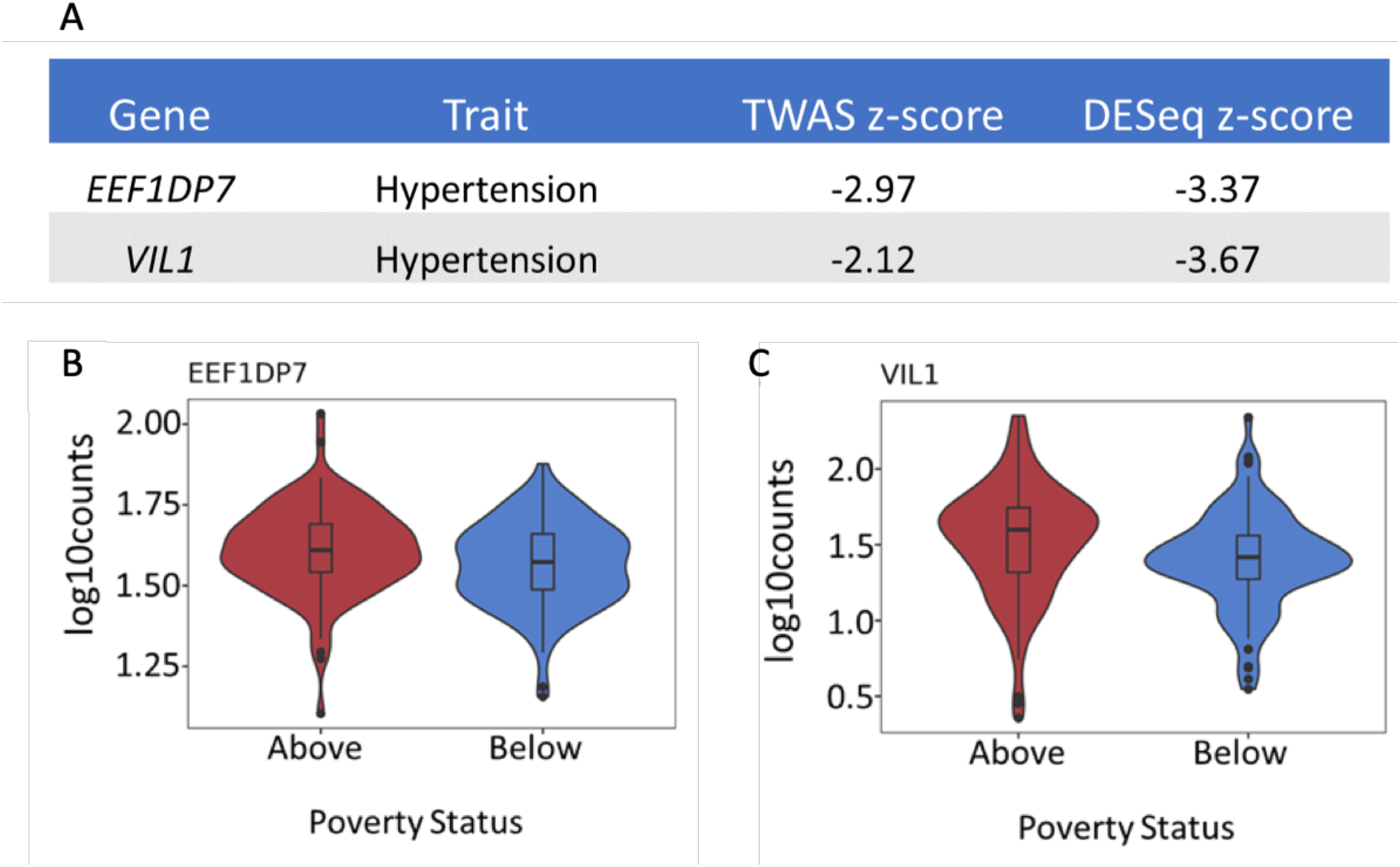
Genes associated with poverty status are also associated with disease risk. A. Table of the differentially expressed genes that are also associated with hypertension in PTWAS. The table columns list the genes, trait, TWAS z-scores, and DESeq z-scores. B-C Violin plots of the two differentially expressed genes associated with hypertension. The x-axis indicates the poverty status, and the y-axis indicates the normalized log10 gene expression counts.

## Discussion

Our study advances the knowledge of how a social determinant of health, poverty, may influence gene expression, using a well-balanced, inner-city cohort. Using RNA-sequencing in a cohort of 204 individuals, we found 138 genes differentially expressed between individuals living either above or below the federal poverty line. We discovered that these genes were enriched for coagulation and wound healing processes. We also found overall stronger effects in women, compared to men. Our results suggest that poverty may contribute to health disparities in hypertension through gene expression changes in immune cells. Our findings identify several important pathways that are differentially regulated by poverty.

Our previously published work conducted on a smaller subset of 52 individuals^22^ found suggestive indication of an association between poverty and immune cell gene expression. Here, using a larger cohort study design and examining genome-wide transcriptional changes, we found significant gene expression changes in individuals living in poverty. Our results show that it is possible to identify transcriptional signatures associated with poverty in well powered population studies. Furthermore, these effects are detected consistently across two time points. Similarly a larger study in a population of young adults^32^, only found suggestive evidence of transcriptional changes associated with poverty. While these changes are largely consistent with our results, we identify stronger effects suggesting that the effects of poverty on gene expression is stronger in older individuals that may have been chronically exposed to poverty.

We identified wound healing among the significant biological processes dysregulated in individuals that live in poverty. The role of immune cells in wound healing is critical for integrity restoration of damaged tissues^54^. Chronic inflammation of immune cells delays tissue wound healing repair^54,55^. Monocytes are one of the cell types in PBMCs and single cell studies have shown that their transcriptional status influences other PBMC components (e.g. T cells)^56,57^. Lack of plasticity of monocytes/macrophages in the inflammatory to anti-inflammatory signaling promotes ineffective wound healing, which is a main complication of type 2 Diabetes^58^. In addition, chronic pro-inflammatory states of monocytes have been linked to cardiovascular diseases like coronary artery disease and myocardial infarction^59,60^. Here we analyzed gene expression within a mixed population of peripheral blood mononuclear cells and did not discern whether transcriptional profiles were associated with unique cell types. Follow-up analyses are warranted to experimentally validate these initial findings and identify if transcriptional profiles in specific cell types serve as better biological biomarkers for the effect of poverty status on gene expression.

Men and women differ in their risk for many complex diseases (e.g.coronary artery disease and systemic lupus erythematosus^61,62^). These sex differences are attributed to hormones, sex chromosomes, genotype-sex effects, behavioral differences, and differential environmental exposures; however, the mechanisms and underlying biology need clarification. Oliva et al^46^, studied differences in gene expression between men and women in the Genotype-Tissue Expression (GTEx) dataset and identified 13,294 genes differentially expressed in at least one tissue. In blood they found 1710 differentially expressed genes. The sex-specific differences in gene expression observed between individuals living in poverty or above poverty may reflect differences between sexes in average life expectancy of at-risk populations^63^. Compared with individuals who are not living in poverty, we did not observe gene expression changes in men living in poverty, but we did find significant changes in women (Figure 2). Gene expression differences between men and women could reflect that living in poverty has a less profound effect on gene expression levels in PBMCs in men.

Poverty is consistently linked to social and environmental conditions that act as stressors and are associated with risk of chronic illness^64^. Earlier studies in social genomics identified a pattern of differentially expressed genes known as the conserved transcriptional response to adversity (CTRA)^16,17,19^. Socioeconomic status and psychosocial experiences have a broader effect on immune cell gene expression and impact additional pathways beyond those included in the CTRA^28^. We identified 138 differentially expressed genes, of which only *PTGS1* was also included in the CTRA. Our study thus identifies novel genes and biological processes and pathways associated with poverty that were previously unknown.

Here we focused on poverty strictly defined as household income, however several factors of living in poverty may be directly associated with gene expression differences, including for example food insecurity or nutritional factors, psychosocial stress or perceived violence, or any other complex environmental influences resulting from living in an impoverished urban area. Future studies aiming to dissect the relative effect of each of these factors will be important to guide interventions aimed at improving health outcomes for inner-city residents.

In conclusion, our study provides evidence that poverty influences gene expression patterns in a large diverse cohort. Impoverished environments have long been associated with the prevalence of poor health outcomes. This work helps elucidate the molecular pathways that potentially transduce the effects of poverty into poor health outcomes, thus providing an understanding of how social determinants of health alter gene expression and can provide additional information about pathways that contribute to disparities in health outcomes in disadvantaged populations.

## Materials and Methods

### Study Participants and sample collection

The Healthy Aging in Neighborhoods of Diversity Across the Life Span (HANDLS) study^65^ is a longitudinal, epidemiological study of a diverse cohort of individuals living in Baltimore City and was designed and implemented by the National Institute on Aging Intramural Research Program (NIA IRP). Poverty status was assigned based on self-reported household income, by grouping participants as living either 125% above or below the 2004 federal poverty line. Race was self-reported as either African American or White. Samples were collected at two different time points ∼4-5 years apart for each participant (time point 1 and time point 2). Participant blood samples were collected in the morning. The isolation of the PBMCs was conducted within 3 hours of blood draw, aliquoted, and stored at -80°C. The Institutional Review Boards of the National Institute of Environmental Health Sciences, National Institutes of Health and Morgan State University approved this study and a written informed consent was signed by all participants.

In the present study, HANDLS participants were selected across 4 variables: sex, race, poverty status, and age group (10-yr groups between 35-65). The cohort was balanced for all variables and divided in 24 batches for a total of 240 individuals across two time points, and 480 samples. Following RNA-sequencing QC (see section below), the final sample size consisted of 204 participants (Figure 1B).

### RNA Extraction, Library preparation and Sequencing

Cells were thawed and transferred into RPMI1640 (Gibco Cat# 11835030) medium containing 10% FBS (Gibco Cat# 10082147). Cells were then collected by centrifugation at 1200 RPM for 7 min and washed twice using ice-cold PBS (Gibco Cat# 10010002). Collected pellets were lysed using Lysis/Binding Buffer (Invitrogen Cat# 61012), and frozen at -80°C. Polyadenylated mRNAs were subsequently isolated from thawed lysates using the Dynabeads mRNA Direct Kit (Invitrogen), following the manufacturer’s instructions. RNA-seq libraries were prepared using a protocol modified from the NEBNext Ultradirectional (Illumina® -NEB Cat# E7760L) library preparation protocol. RNA was fragmented at 94°C for 5 minutes to obtain 200-1500 bp fragments. SPRISelect beads (Beckman Coulter) were used in all purification steps and size selection. Barcodes from PerkinElmer were added by ligation. The individual libraries were quantified using the KAPA real-time PCR system (ROCHE DIAGNOSTICS Cat # 07960441001), following the manufacturer’s instructions and using a custom-made series of standards obtained from serial dilutions of the PhiX DNA (Illumina). Next, libraries were pooled and sequenced using the Novaseq instrument. Median number of reads/sample was 33,015,582 Supplementary Table 1.

### Data pre-processing

We aligned the RNA-seq reads with HISAT2^66^ to the human genome version ‘grch38_snp_tran’, which considers splicing and common genetic variants. Aligned and cleaned (deduplicated) reads were counted using HTSeq and Homo_sapiens.GRCh38.103 (Ensembl) transcriptome assembly genes. We started with an initial sample size of 478 libraries corresponding to 239 individuals. Due to poor RNA quality 26 libraries failed to sequence, which resulted in 452 libraries. For all gene expression analyses, genes on sex chromosomes and genes with expression below six reads or 0.1 counts per million in at least 20% of samples were dropped, which resulted in 21,251 genes.

To validate the identity of the two time point samples from the same individual, we first used the program ANGSD ^67^ to infer genotype likelihoods from the aligned and cleaned mapped reads. The program NgsRelate^68^ was then used on the genotype likelihoods to estimate relatedness between all sample pairs. Individual sample pairs from the two time points were retained if the pair had an estimated KING coefficient greater than 0.30. This resulted in a final set of 204 individuals (408 samples) that were used in all subsequent analyses.

We obtained genotype information from the RNA-seq data by using samtools pileup and bcftools on known genotypes from the 1000 genomes project^69^. The resulting VCF file was filtered for biallelic SNPs and the samples were further analyzed with PLINK2^70^. The first 3 genotype principal components were added in the gene expression model to control for genetic ancestry. Additionally, the following covariates were included in the model to correct for confounders: library preparation batch, age, sex, percent reads mapping to exons, percent non-duplicate reads.

### Differential gene expression analysis

We used DESeq2 v1.22.1^71^ to test for differential gene expression across 204 individuals in each time point.

Model 1 (Poverty Model): library preparation batch + age + sex + percent reads mapping to exons + percent non-duplicate reads + PC1 + PC2 + PC3 + poverty status

We also examined each sex separately:

Model 2 (Women and Men Split Poverty Model): library preparation batch + age + percent reads mapping to exons + percent non-duplicate reads + PC1 + PC2 + PC3 + poverty status

To control for FDR, we used the Benjamin-Hochberg independent filtering step and multiple test correction implemented in DESeq2. Supplementary Table 2 contains the full results of the analysis.

### Gene Ontology and pathway enrichment analyses

To identify relevant pathways related to the significantly differentially expressed genes in individuals living in poverty, we performed an enrichment analysis (hypergeometric test) for GO, KEGG and REACTOME terms with clusterProfiler in R^72^. We tested each list of differentially expressed genes against a background of all tested genes in our dataset. Significant biological processes contained a minimum of 10 genes and a maximum of 500 genes with a p adjusted < 0.1.

### TWAS analyses

To directly investigate whether discovered effects on gene expression may contribute to differential health outcomes we used the results from PTWAS (Probabilistic Transcriptome-Wide Association Study)^47^ (5% FDR) for cardiovascular disease traits such as: Coronary Artery Disease and hypertension. We overlapped the significantly differentially expressed genes associated with poverty and focused on those genes that had a z-score larger than 2 in absolute value^47^.

## Data Availability Statement

All RNA-seq files are available from the Gene Expression Omnibus database (accession number GSE221766).

## Acknowledgments

We thank Samuele Zilioli and members of the Luca/Pique-Regi group for helpful discussions and comments on an early version of this manuscript. We would like to thank the HANDLS participants and clinical staff, and Nicolle Mode for helping with cohort design. This work was supported by the NIH grant #U54MD013376 and by the National Institute on Aging Intramural Research Program, National Institutes of Health project #AG000513.

## Supplementary Information

Supplementary Table 1

Number of reads for each of the 408 libraries. Listed in three columns, the libraries, reads, and the non-duplicated reads.

Supplementary Table 2

DESeq 2 results from model 1, includes the base mean of the gene, the log2 fold change, standard error, z-score, p-value, and the adjusted p-value for all genes tested.

Supplementary Table 3

Results of GO over-representation analysis, columns include: gene ontology term ID, description of the term, gene ratio, background ratio, p-value, adjusted p-value, q-value, gene ID, and the counts of genes associated to each term.

**Supplementary Figure 1:**
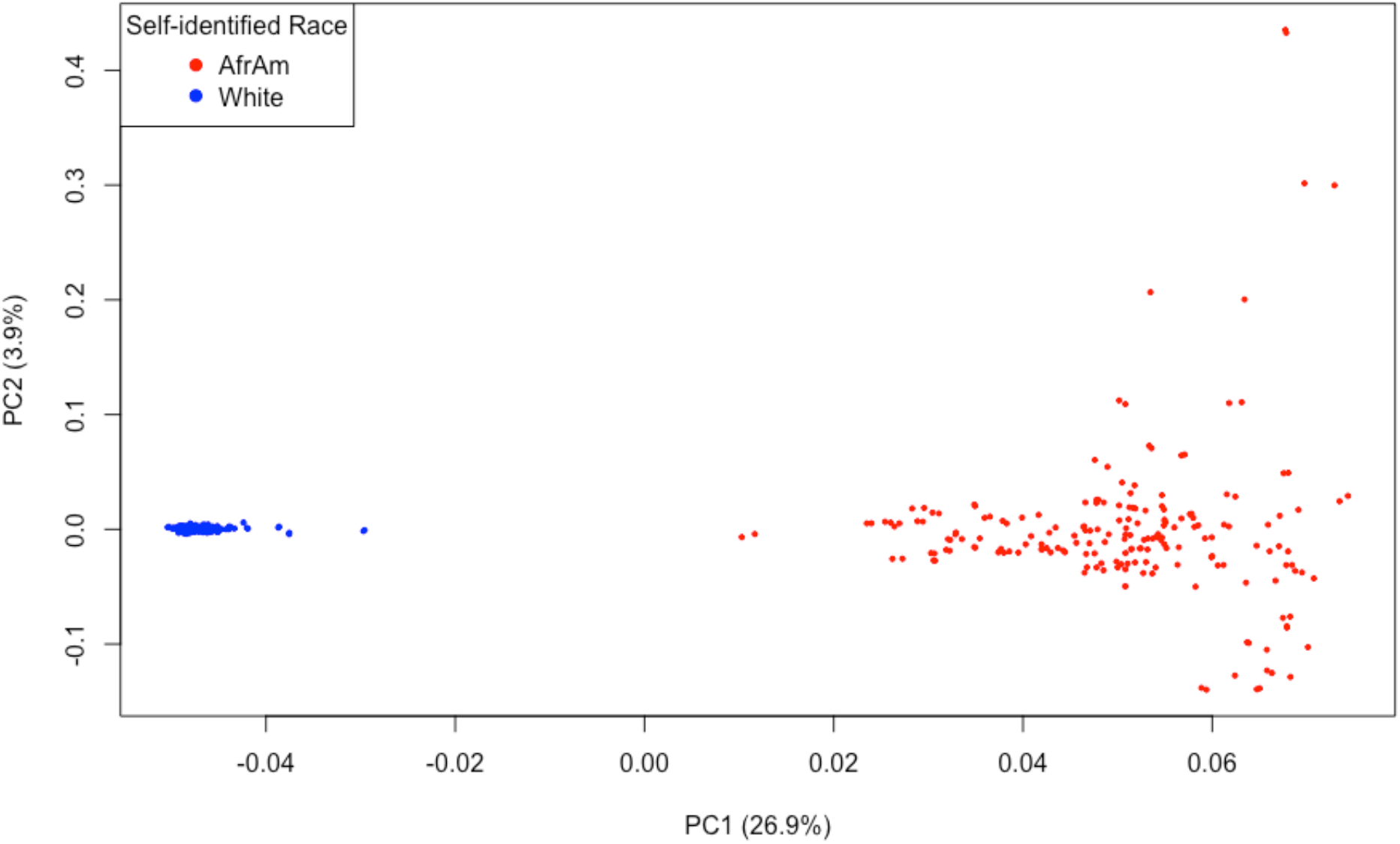
Principal Component Analysis of individual genotypes. Scatter plot of principal component 1 (x axis) and principal component 2 (y axis) calculated on the genotypes inferred from the RNAseq reads. Each dot represents an individual and their self-identified race (African American in red and White in blue).

**Supplementary Figure 2:**
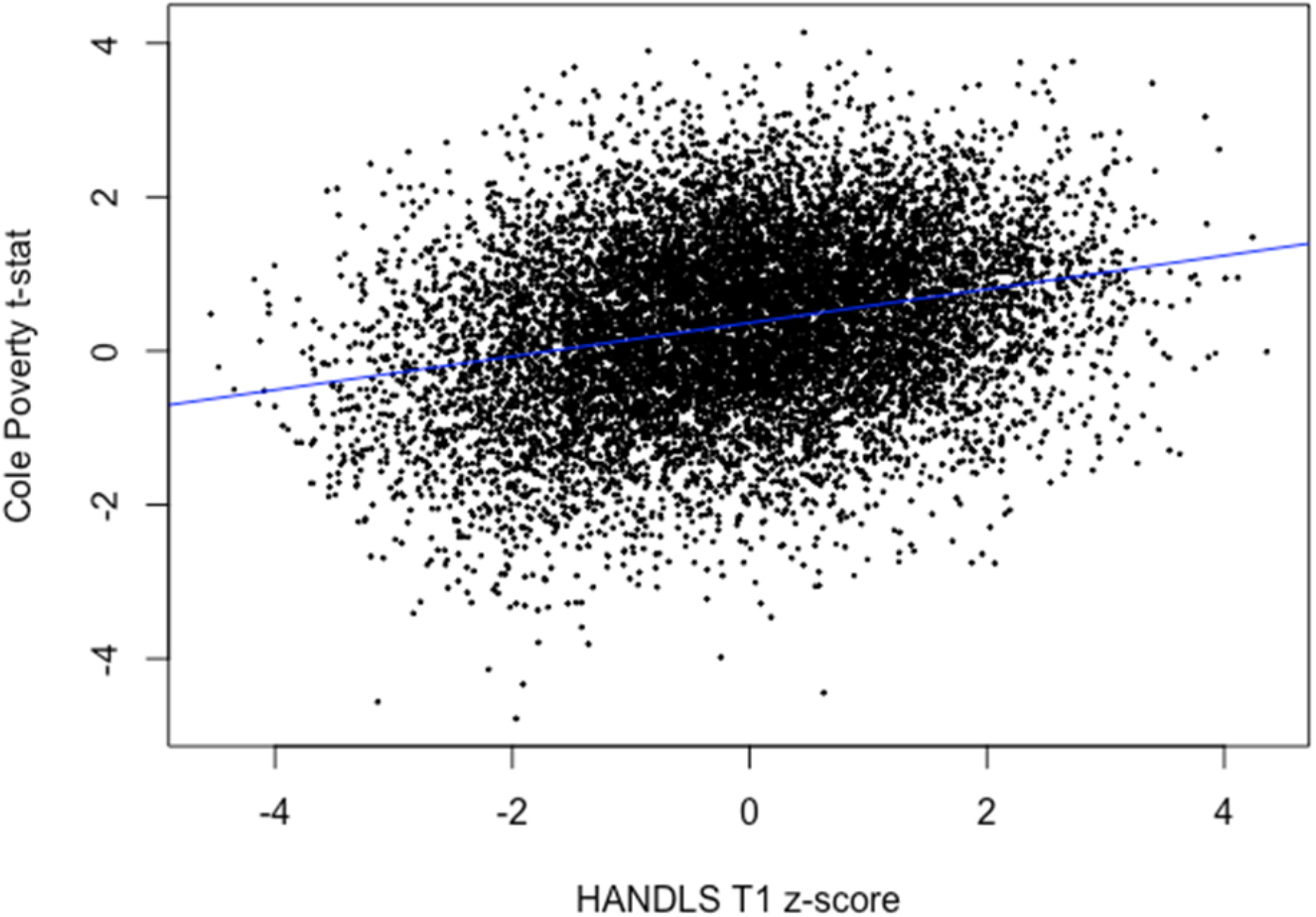
Scatter plot of z-scores for poverty effects in the current study and the Add Health study^32^. Each point represents a gene. The blue line is the regression line (ρ=0.264 and p-value< 2.2e-16).

